# Symbolic labelling in 5-month-old human infants

**DOI:** 10.1101/414599

**Authors:** Claire Kabdebon, Ghislaine Dehaene-Lambertz

## Abstract

Humans naturally entertain complex representations of the world based on various symbolic systems, from natural language to mathematical or musical notation. Above and beyond mere perceptual representations, the adult human mind can recode sensory inputs into abstract symbolic representations that can be internally manipulated and projected back onto the external world. However, the ontogeny of this striking ability remains controversial: Are children progressively acquiring symbolic representations through language acquisition, or are mental representations symbolic from the very beginning, language learning consisting in mapping mental symbols onto public symbols? Using high-density electroencephalography, we show here that preverbal infants can form mental representations that feature symbolic attributes. In three experiments, a total of 150 five month-olds were exposed to triplet words characterized by their abstract syllabic structure (AAB/ABA/ABB) consistently followed by an arbitrary label. Subsequently, incongruent structure-label pairings evoked a late violation-of-expectations signal, whereas congruent pairings induced an early priming effect. Importantly, the late surprise response was recorded for incongruent pairs even when the pairing order was reversed at test (i.e. labels preceding structure). Our results indicate that first, far beyond habituation/dishabituation, preverbal infants are able to recode sensory inputs, into abstract mental representations to which arbitrary labels can be flexibly assigned. Second, we demonstrate that, beyond conditioned associations, this mapping is instantly bidirectional. These findings buttress the hypothesis of symbolic representations in preverbal infants, which may serve as a foundation for our distinctively human learning abilities.

**Significance Statement:** Symbolic systems provide a powerful tool for efficiently re-describing the world into operable mental variables that, in turn, become objects of cognitive manipulation. However, is this ability tied to mastering language? Using an associative learning task in preverbal infants, we show that 5 month-olds can re-describe percepts into abstract mental variables that can be associated with arbitrary labels, well before they produce their first words. Importantly, we show that, beyond associative learning, they readily inferred a bidirectional mapping between the re-described representations and the associated labels, a capacity that animals do not spontaneously exhibit. Human cognitive success might be rooted in such abstract recoding which is no longer sensitive to local variations, thus alleviating cognitive load, and ultimately facilitating learning.

---

Human intelligence consists in representing the world way beyond the direct evidence of our senses. From the first months of life, human infants have access to more than perceptual features, as they are sensitive to several abstract attributes of their environment (1). However, the initial format of early mental representations remains unclear. On the one hand, proponents of the Quinian perspective (2) propose that infants progressively transition from a state where information is implicitly encoded as distributed representations, hidden in a mesh of cerebral connections, to a state where mental representations are explicitly represented as linguistic variables thanks to the progressive acquisition of the symbols and rules of natural language. This hypothesis is supported by animal studies indicating that non-verbal primates exhibit abstract reasoning abilities when they are trained to use an explicit symbol system (3-5). However, the extensive training required for non-human primate to acquire the provided symbols, together with their rather imprecise behavioral outcomes tends to suggest that the abstracted mental representations still radically differ from that of human adults. Rationalists (6), on the other hand, argue that, from start, the human mind readily recodes perceptual input into localized abstract mental variables (or mental symbols) which support a subsequent referential mapping with the public linguistic symbols used in the society during language acquisition.

The study of abstract knowledge in preverbal infants focused on whether, or not, they could encode abstract features based on habituation/dishabituation procedures. For example, 7 month-old infants (7-10) and even newborns (11) habituate to the presentation of short auditory or visual sequences based on their repetitive structure, and at test they are able to discriminate between familiar and novel structures. Yet, the question of the encoding format of these representations remains unaddressed. Both distributed and localized mental representations can translate into successful discrimination abilities. However, only information encoded as a localized mental representation can be manipulated and passed along to further processing stages. Given this assumption, if the preverbal mind is limited to capturing abstract information as distributed representations, it should not be able to operate over abstract knowledge. On the contrary, if preverbal infants can recode perceptual input as abstract mental variables, they should be able to manipulate abstract knowledge, and concurrently entertain multiple representations that can support subsequent mapping with an arbitrary artefact. In the present study, we tested this hypothesis, asking whether preverbal infants could manipulate abstract representations, hence exploring the symbolic attributes of preverbal mental representations.

We used high-density electroencephalography in three associative learning experiments (conducted with 48, 57 and 45 infants) in order to investigate the mental representations of 5-month-olds. Participants were presented with tri-syllabic nonce words characterized by their repetition-based structure (10) (AAB/ABA/ABB), followed 1 second later by the presentation of an arbitrary label. Crucially, each abstract structure was paired with a given label. Under these conditions, infants could not rely on the mere perceptual input but had to encode the underlying abstract structures as operable mental variables in order to categorize the different words and discover the systematic pairings.

## Results and discussion

In experiment 1, after a short exposure to AAB and ABA words paired with arbitrary images (Figure 1-A), learning was subsequently assessed during a testing phase, when we introduced a small proportion of incongruent pairings (25%) and a novel ABB structure, which was equally paired with both images (non-predictive condition). Overall for each infant, both images were presented with equal frequency. If infants successfully represent the abstract word structures and detect the associations, we would then expect to record different brain signatures in response to the congruent and incongruent images. Non-parametric analyses of the event-related potentials (ERPs) revealed a late negative activation in response to incongruent images compared to congruent images (the late slow wave, LSW), yielding a significant negative difference on a cluster of right central electrodes (Monte-Carlo *p* = 0.036 in 32 infants, Figure 2.A). Further comparisons revealed that brain responses to images following the ABB structure, which was not predictive of any image, remained close to the congruent condition (*t*(31) = 1.06, *p*_*corr*_ = 0.597) but significantly different from the incongruent condition (*t* 31 = −2.84, *p*_*corr*_ = 0.016), indicating that this late activation reflected sensitivity to a violation-of-expectations^5^. This result is in line with previous results suggesting that the LSW recorded in infants uncovers similar but much slower processes as the adult P300 component (12).

**Fig. 1:**
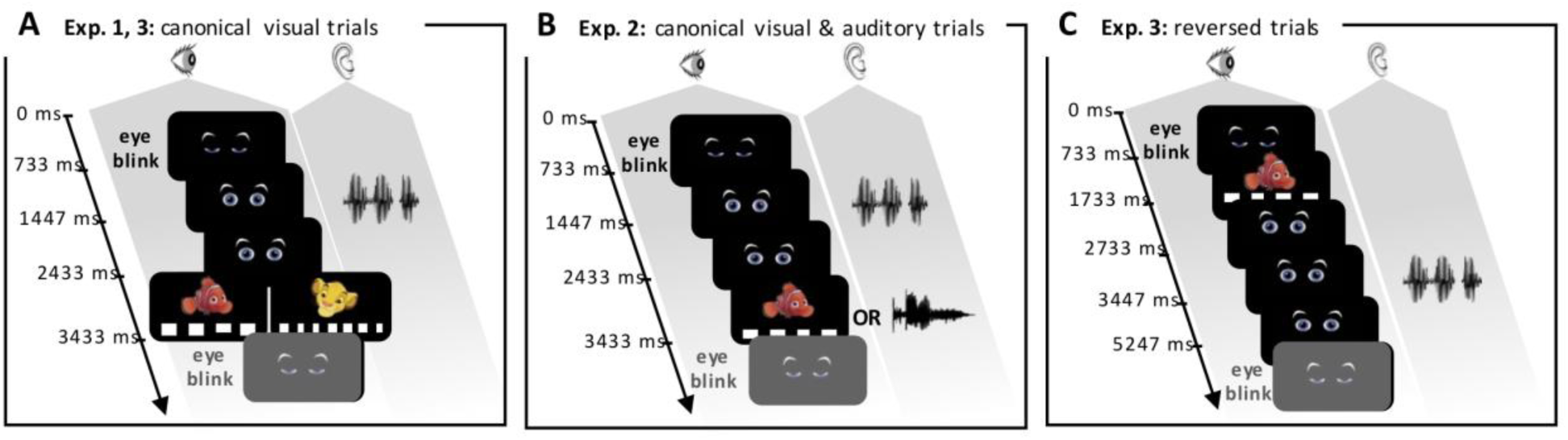
Experimental paradigms. Trials began with blinking eyes. In canonical trials, a tri-syllabic word was then presented, followed approximately 1s later by the presentation of an image (a lion or a fish) presented on a flickering background (10 and 15 Hz) in experiments 1 and 3 (A), and a fish or a word (“*Schtroumpf*”) in experiment 2 (B). Infants first learned the association between two structures and their labels during a 36-trial exposure phase (e.g. ABA-lion and AAB-fish). During the subsequent test phase, incongruent structure-label pairs (e.g. ABA-fish and AAB-lion) and a new structure (e.g. ABB-fish and ABB-lion) were introduced in experiments 1 and 2. In experiment 3, the test phase consisted of an alternation of short blocks of 6 reversed trials (C) and longer blocks of 14 canonical trials (A). Canonical trials were all congruent contrary to reversed trials, for which half were congruent and the other half incongruent.

**Fig. 2:**
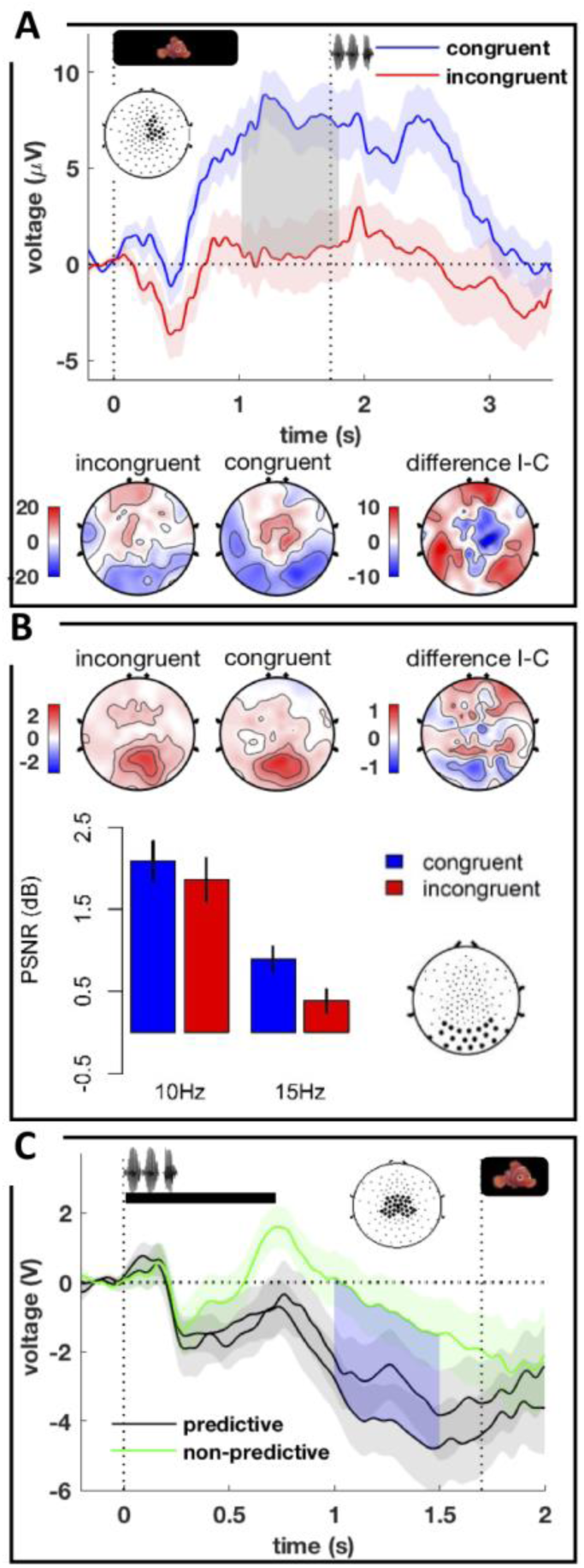
Three neural signatures of learning (experiment 1). A-Late violation of expectations. Top row: grand average for congruent (blue) and incongruent (red) trials recorded from the significant cluster of electrodes presented on the graph. The vertical dotted line at 1.733s indicates the onset of the next trial. Bottom row: ERP topographies for incongruent and congruent trials as well as their difference averaged over the significant time-window (grey area on the plot). Electrodes and time were identified using non-parametric analyses. **B - Early priming effect**. Top row: PSNR topographies in response to the flickering background during label presentation for both conditions and their difference. Bottom row: bars represent PSNR averaged over the occipital cluster in response to congruent (blue) and incongruent (red) trials for 10Hz and 15Hz backgrounds. PSNR was significantly larger for congruent compared to incongruent trials**. C - The Contingent Negative Variation**. Grand average for the two predictive structures (black) and the non-predictive structure (green), recorded from central electrodes. The blue area indicates the significant time window and the vertical dotted line at 1.700s, the onset of the visual label. The predictive structures (AAB & ABA) elicited a significantly larger CNV than the non-predictive structure.

A careful inspection of the trials comprising ABB words allowed us to ask whether infants were representing the entire tri-syllabic sequence, or whether they were only detecting immediate repetitions. In the latter hypothesis, they should similarly consider <u>AA</u>B and A<u>BB</u> structures and thus expect the same image in both cases. In terms of EEG, this assumption implies a violation-of-expectations when the presented image after an A<u>BB</u> word is not the image they learned to follow <u>AA</u>B words. On the contrary, according to the former assumption, they should successfully entertain all three structures as distinct mental representations and realize that ABB words are not predictive of any specific image. When contrasting brain responses to the two images following ABB words, we found no violation of expectations (*t*(31) < 1), indicating that infants’ expectation of an image was indeed based on the full description of the abstract relations between all three syllables in the word.

Beyond late violation-of-expectations, we wondered whether the expectations that infants derived from the abstracted structures might also prime early visual activity for receiving a precise input (13, 14). Because each image was presented over a background flickering at a specific frequency, we inspected whether the strength of the entrained low-level visual activity was modulated by the congruency of the structure-image pair on a cluster of 20 occipito-temporal electrodes commonly used in the literature to study steady-state responses (15). A significant enhancement of cortical entrainment was observed for the expected frequency compared to the unexpected frequency (*F*(1,31) = 7.97 *p* = 0.008; Figure 2.B). Furthermore, when observing brain responses prior to image onset, we observed a negative deflection over central electrodes which developed after the end of the triplet. This waveform corresponds to the Contingent Negative Variation (CNV), commonly recorded after a cue predictive of a target stimulus in adults (16) as well as in infants (17). Its amplitude was affected by word structure (39 infants, main effect of structure: *F*(2,76)=4.98, *p* = 0.009 Figure 2.C): it was larger after the AAB and ABA words that were predictive of the forthcoming image than after the non-predictive ABB words (AAB vs. ABB: *t*(38) =-2.03, *p* = 0.049, ABA vs. ABB: (*t* (3)= −2.63, *p* = 0.012), whereas there was no difference between AAB and ABA trials (*t*(38)= −1.41, *p* = 0.168).

Altogether, these results demonstrate that infants successfully represented all three structures concurrently, without confusion, and in a format that allowed for a subsequent, immediate association with an arbitrary image. For infants to succeed in our experimental design, it was not sufficient to locally discriminate between two structures as in classical habituation-dishabituation paradigms; rather, infants had to have access to an abstract summary of the sequence of syllable repetitions and changes in order to couple it with the subsequent image. In other words, they had to re-describe the auditory input into one mental structure that could be further manipulated. The predictive activity observed during the waiting period between the offset of the tri-syllabic word and the onset of the image may reflect this manipulation stage offering infants the possibility to prospectively infer the associated image. Overall, experiment 1 demonstrates that preverbal infants form mental representations that are not limited to the output of perceptual processes but rather involve a representational re-description stage (18), which makes knowledge available to the mind for other mental operations, a fundamental feature of symbolic representations. In experiments 2&3, we inspected two aspects of the relation between the generated mental representations and the arbitrary labels: flexibility and bidirectionality.

Since human adults are remarkably flexible in associating any mental representation with virtually any type of symbol (words, numbers, graphs, pictures, shapes, etc.), we asked, in experiment 2, whether 5-month-old infants were just as versatile and might accept labels in different sensory modalities (Figure 1.B). The procedure was the same as in experiment 1 except that the lion image was replaced by an auditory word. In both auditory and visual modalities, a significant violation-of-expectations signal was recorded over right central clusters peaking after 1 second post label onset (for the visual label: Monte-Carlo *p* = 0.038 in 34 infants; for the auditory label: Monte-Carlo *p* = 0.0048 in 32 infants; Figure 3). Combining visual responses in experiments 1 and 2, non-parametric statistics confirmed the common violation-of-expectations response over a right central cluster (Monte-Carlo *p* = 0.024 in 66 infants). We thus replicated the late surprise effect induced by an incongruent pairing, even when the label modalities were mixed. Like toddlers, who accept any arbitrary form (gestures, pictograms, or sounds) as a label if embedded in a naming routine (19), infants were similarly able to flexibly associate structures with labels in multiple modalities, suggesting that any artifact, provided it consistently followed the triplet, could be used to represent the given structure.

**Fig. 3:**
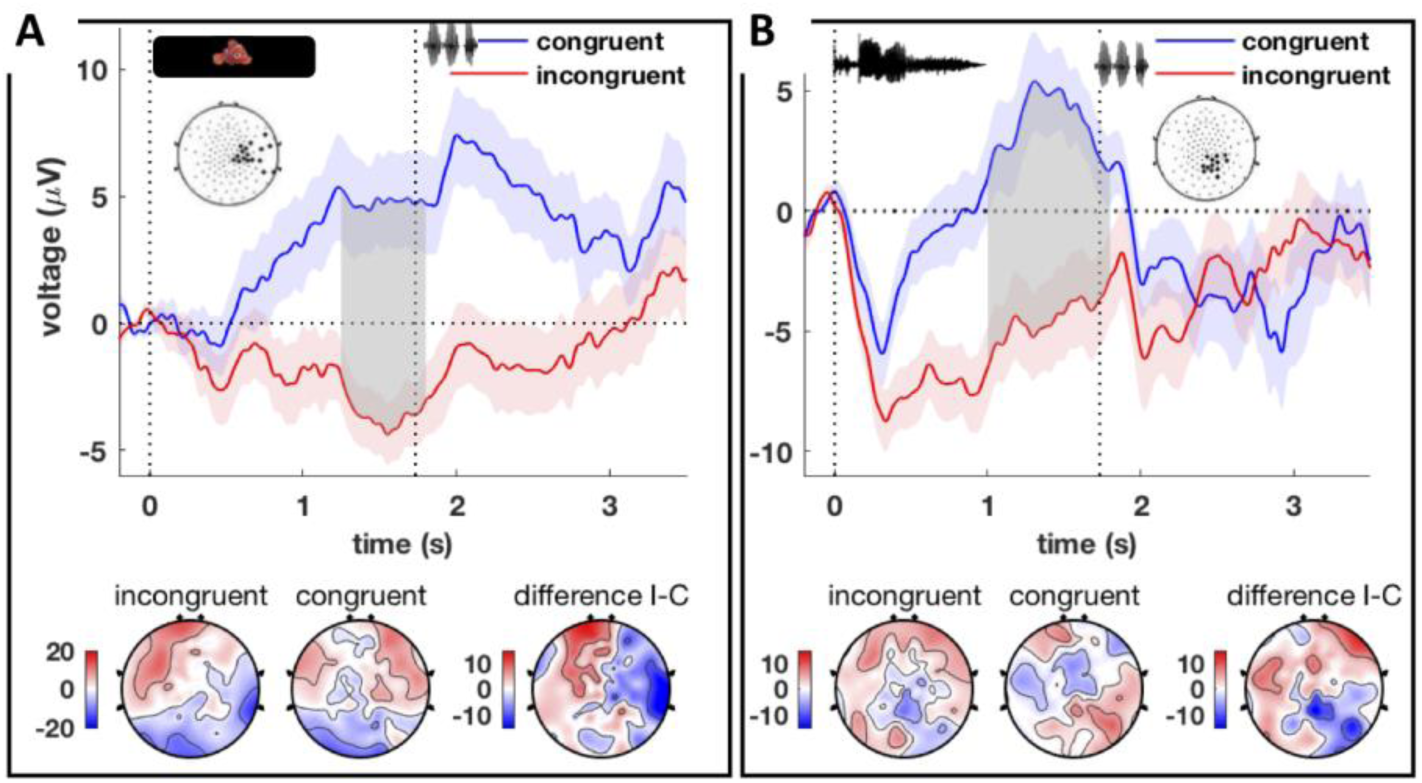
Violation of expectations (experiment 2). Top row: grand average for congruent (blue) and incongruent (red) trials recorded from the significant clusters of electrodes presented on the graphs, for visual (A) and auditory labels (B). The vertical dotted line at 1.733s indicates the onset of the tri-syllabic word of the following trial. Bottom row: ERP topographies for incongruent and congruent trials as well as their difference averaged over the significant time-windows (gray rectangles on the plots) for visual (A) and auditory labels (B). Electrodes and time were identified using non-parametric analyses.

Unlike conditioned associations in which the temporal ordering between the unconditioned stimulus and the conditioned stimulus is crucial, symbolic mapping entails a relation of symmetry, supporting bidirectional predictions between the symbol and the representation to which it is tied. The ability to appreciate bidirectional mappings is readily observed in both human adults and children but appears to be particularly challenging for non-verbal species (20, 21). In experiment 3, we tested whether pre-verbal infants spontaneously inferred a bidirectional pairing between the abstracted structures and their associated arbitrary labels. Infants were presented, after the exposure phase, with short blocks of reversed trials (image-word) embedded within long blocks of congruent canonical trials (word-image). Half of the reversed trials were congruent, and the other half incongruent, preventing any associative learning within reversed trials. If infants possess a symbolic mapping mechanism that can operate over abstract representations, they should spontaneously transfer the structure-image association from canonical to reversed trials, and we would then expect different brain responses to congruent and incongruent trials. ERPs indeed showed a significant congruency effect over a late central dipolar component culminating around 1.7s post word onset (Monte-Carlo *p* = 0.044 in 34 infants; Figure 4). Therefore, not only did 5-month-olds expect to see the appropriate image after having recovered the abstract word structure in canonical blocks, but they also expected to experience the appropriate structure after seeing the image in reversed blocks, without any additional training.

**Fig. 4:**
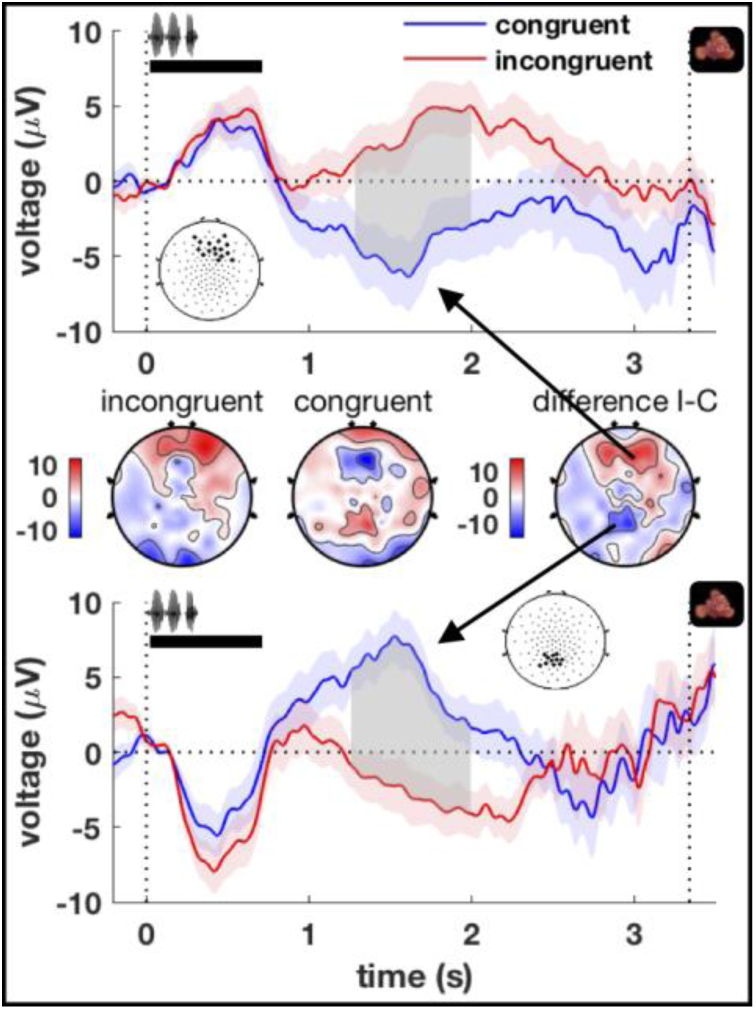
Reversed trials in experiment 3. Top and bottom rows: Grand average for congruent (blue) and incongruent (red) trials recorded from the positive (top) and negative (bottom) significant clusters of electrodes presented on the graph. Middle row: ERP topographies for both conditions and their difference over the significant time-window (gray rectangle on the plots). Electrodes and time were identified using non-parametric analyses. Infants were able to generalize the structure-label pairings from the trained canonical order to the untrained reverse order, demonstrating a bidirectional mapping between the auditory structures and the arbitrary visual labels. The vertical dotted line at 3.247 s indicates the onset of the following trial.

This series of three experiments highlights the symbolic depth of mental representations in preverbal infants, with two important findings. First, we show that well before they begin to speak, infants can combine a series of abstract features (here a sequence of syllable repetitions and changes) into an operable abstract variable, available for subsequent associative operations. While classical habituation/dishabituation paradigms merely address information encoded *in* the infant mind, as distributed implicit network states, our paradigm allows to pinpoint abstract, re-described representations that are available *to* the infant mind. Contrary to previous speculations that infants merely use labels as mnemonic cues (22), attentional grabbers (23) or essence placeholders (24), we propose here that labels drive infants into representational re-description and help them organize their inner mental space through a mapping between abstract representations and their associated labels. Importantly, these results show that language proficiency is not a prerequisite for combining different abstract features (here, the presence of a repetition and its ordinal position) into a unitary mental representation.

Second, we demonstrate that these preverbal yet sophisticated mental representations are flexibly associated with arbitrary labels in multiple modalities and more importantly through a powerful bidirectional mapping. Whereas non-human animals fail to spontaneously generalize conditioned associations to the reverse order (20, 21, 25, 26), humans are particularly prone to disregarding temporal orderings and inferring symmetry (20), sometimes at the expense of logical reasoning (27). It was therefore commonly acknowledged that language experience supported symmetry. Our study, which reports preverbal abilities for symmetry, questions this assumption. Together with previous studies showing that even primates with previous linguistic experience fail to infer bidirectional mappings when learning symbols using conditioned associations (28), this empirical finding suggests that symmetry might be the early marker of an inner representational system, prone to attributing a mental variable to an external category, in other words, reducing it to a sign.

With intensive training, non-human animals (29, 30) also come to use arbitrary labels to represent objects, colors or quantities. Monkeys may even learn to combine abstract signs in formal operations such as addition (3). However, infant behavior, which rapidly and spontaneously infers a powerful bidirectional mapping between a set of abstract representations and a set of labels together with the abundant evidence that non-human animals have difficulties in processing bidirectional associations, suggests a clear discontinuity between human and non-human behavior, which might be a building block for human cognition and notably language development.

## Methods

All experiments were approved by the regional committee for biomedical research. Both parents were informed and provided their written consent before the experiment.

### Stimuli

The tri-syllabic words (duration: 720ms) were generated as the concatenation of synthesized CV syllables to conform to AAB, ABA, and ABB structures. Two different sets of 15 CV syllables were used to build two separate vocabularies of 120 distinct words. One vocabulary was made from the consonants [‘b’, ‘t’, ‘k’] and vowels [‘a’, ‘u’, ‘ou’, ‘in’, ‘e’] and the second vocabulary was made from consonants [‘p’, ‘d’, ‘g’] and vowels [‘an’, ‘eu’, ‘i’, ‘o’, ‘on’]. The syllables were generated with a duration fixed at 240 ms and flat intonation using the MBROLA text-to-speech software (31) with French diphones and digitized at 22050Hz. The syllables were concatenated to form tri-syllabic words (duration: 720 ms), ensuring that the consonant and vowel from syllable A were systematically different from consonant and vowel from syllable B. The labels consisted of two cartoon pictures in Experiments 1&3, and one cartoon picture and one auditory 1-second-long word in Experiment 2.

### Protocol

Brain activity was recorded using a high-density EEG net (128 channels, EGI, Eugene, USA) while infants were seated on their caregivers’ laps. A typical canonical trial consisted in the presentation of a 720 ms auditory word, followed by a 1 second silence delay before the onset of one of the two labels. We used the image of a red fish and the image of a yellow lion as the two visual labels, and each image was systematically presented over a background flickering at a specific frequency (10Hz or 15Hz) during 1 second. The auditory label consisted of a 1 second monosyllabic word “*Schtroumpf*” recorded from a female native French speaker in an infant-directed speech register. During the exposure phase, infants were presented with a series of 36 of such trials in which the two labelled structures were consistently paired with their respective labels. The pairings were counterbalanced across participants.

#### Experiment 1

47 healthy 5-month-old infants were recruited to participate: 15 participants were excluded from analyses of the visual labels (final group: 14/18 girls/boys, 21 weeks +/-2 weeks, range 17-26 weeks), and 8 infants were excluded from analyses of the tri-syllabic words (final group: 16/23 girls/boys, 21 weeks +/-2 weeks, range 17-26 weeks). After the exposure phase in which infants were introduced with AAB and ABA words paired with the red fish and the yellow lion, we introduced certain variations: First, tri-syllabic words conformed to either the two exposure structures AAB and ABA, or to a third structure ABB, which was equally paired with the two images. Second, in 25% of AAB and ABA trials, the visual labels were swapped such that the associations were incongruent with the learned pairings. Overall, the two visual labels as well as the three structures were presented with equal frequency. In this experiment, the generalization vocabulary presented during testing differed from that used during exposure.

#### Experiment 2

57 healthy 5-month-old infants were recruited. The visual and auditory labels were analyzed separately. We report on 34 infants for the visual label analyses (20/14 girls/boys, 20 weeks +/-2 weeks, range 18-24 weeks), and 32 infants for the auditory label analyses (19/13 girls/boys, 20 weeks +/-2 weeks, range 18-24 weeks). Experiment 2 was built on the same model as Experiment 1, except that infants were initially introduced with ABB and ABA words paired with the image of the red fish and the auditory label “*Schtroumpf*”, and AAB was the third structure, equally paired with both visual and auditory labels. In this experiment, both vocabularies were used during testing, and we set aside the generalization vocabulary for a subset of congruent, incongruent and AAB trials used for the statistical analyses.

#### Experiment 3

44 healthy 5-month-old infants were recruited to participate: 10 participants were excluded from subsequent analyses (final group: 11/23 girls/boys, 20 weeks +/-1 week, range 18-23 weeks). Infants were first exposed to ABB and ABA words, paired with the red fish and the yellow lion, respectively, in the canonical structure-label order. During testing, small blocks of 6 reversed trials, were interspersed in between larger blocks of 14 canonical trials, always congruent. In reversed trials, the image was presented first during 1 second, followed by a 1-second silence period and finally a tri-syllabic word was presented, conforming to either an ABB or an ABA structure. Canonical trials systematically conformed to the congruent structure-label pairings. On the contrary, half of the reversed trials were incongruent with the learned associations, so that infants could not learn any label-structure associations from the reversed trials. The generalization vocabulary was only used in the reversed blocks.

### Event related potentials

The electroencephalogram (EEG) was continuously digitized at 250 Hz from 128 scalp electrodes referenced to the vertex. For each channel, we rejected epochs with fast average amplitude exceeding 250µV, or when deviations between fast and slow running averages exceeded 150µV. Participants with less than 10 trials in one of the experimental conditions were rejected from subsequent analyses. The remaining trials were averaged in synchrony with stimulus onset (either labels or tri-syllabic words, depending on the analysis), digitally transformed to an average reference, band-pass filtered (0.2 – 15 Hz), and corrected for baseline over a 200-ms window before stimulus onset (similar results were observed with the raw, unfiltered data), or over the first 200-ms post tri-syllabic word onset in experiment 3, where we could capitalize on the signal alignment, which was easier with the sharp auditory response.

#### Congruency effects

In experiments 1 & 2, given that the proportion of congruent and incongruent trials was strongly imbalanced (75% vs 25% respectively), a subset of congruent trials was defined in order to match the number of incongruent trials. In experiment 1, for each incongruent trial, we selected the closest congruent trial, controlling for the identity of the image (e.g., for each fish-incongruent trial, we selected the closest fish-congruent trial) to control for slow drifts (22 artifact-free trials per subject in the Congruent/Incongruent conditions on average). In experiment 2, the subset was defined upstream (15/16 artifact-free trials on average per subject in the Congruent/Incongruent visual conditions; 16/16 artifact-free trials per subject in the Congruent/Incongruent auditory conditions on average). In experiment 3, the proportion of incongruent and congruent trials was perfectly balanced (16/15 artifact-free congruent/incongruent trials per subject on average). Brain responses to congruent and incongruent trials were systematically compared using non-parametric statistics, combining a clustering and randomization procedure (32), with an alpha threshold set to 0.1, a minimal cluster size of 3 electrodes, and 5000 permutations. This procedure allowed for the detection of positive, negative, and dipolar components (33). Late congruency effects in response to the visual and auditory labels were inspected over a 1-1.8s time window in experiments 1&2, and congruency effects in response to the tri-syllabic words were inspected over a 1-2s time window in experiment 3.

#### Contingent Negative variation

Because the contingent negative variation typically develops between the offset of a cueing stimulus and the onset of the following target stimulus over central recording sites (16), we inspected brain responses during the silence period before label onset in experiment 1 and defined a central cluster of electrodes (32 electrodes around Cz) based on the main response with all trials averaged together. Brain activity was averaged over this cluster for each condition, between 1s and 1.5s post-word onset and was submitted to a three-way ANOVA with Structure (AAB, ABA or ABB) as a within-subject factor. On average, we report on 42 artifact-free trials per subject and per condition (41/42/42 for AAB, ABA, and ABB, respectively).

### Frequency tagging

Brain signals were digitally transformed to an average reference, band-pass filtered (0.2 – 40 Hz), epoched around label presentation, and artifacts were rejected using the same procedure as for ERPs. On average we report on 22 artifact-free trials per subject in the Congruent/Incongruent conditions. Entrainment was measured over an occipital cluster of electrodes (20 electrodes encompassing electrodes Oz, T5 and T6), during picture presentation delayed by 200 ms to compensate for the conduction time of the stimulation to the visual cortices (i.e. 200-1200 ms after stimulus onset). For each subject, each electrode and each trial, we computed the power spectral density (PSD) of the signal over this time-window, using the fast Fourier transform algorithm as implemented in MATLAB. We thereafter averaged the PSDs across trials, for each condition of interest separately for each stimulation frequency (10Hz and 15Hz). From the PSD, entrained activity was measured as the strength of the peak of oscillatory power elicited by the flickering background relative to the local fluctuations of oscillatory power in the surrounding frequencies (Peak Signal-to-noise Ratio or PSNR). Because the power spectrum of brain activity typically conforms to a power-law function, we first fit each average PSD with a power law function (i.e. in the log-space, an affine function using MATLAB polyfit). The PNSR was then computed as the ratio between the deviation of the signal power from power-law at tag frequency, and the root-mean-squared deviation from the power-law measured over neighboring frequencies, in decibels. We averaged the normalized power measurements over the cluster and submitted these values to a univariate analysis of variance with condition (Congruent vs Incongruent) and stimulation frequency (10Hz vs 15Hz) as within-subject factors.

## Acknowledgements

This research was supported by grants from the Fondation Bettencourt-Schueller and ERC AdvGrants Babylearn. The authors would like to thank all the infants and their parents who participated in this study as well as Antoinette Jobert, Giovanna Santoro, the medical team of UNIACT at Neurospin and the LSCP BabyLab in Paris, who helped in carrying out the experiments.

## References

1. Carey S (2009) The origin of concepts (Oxford University Press).

2. Quine WVO (1960) Word and object (MIT press).

3. Livingstone MS, et al. (2014) Symbol addition by monkeys provides evidence for normalized quantity coding. Proceedings of the National Academy of Sciences 111(18):6822–6827.

4. Livingstone MS, Srihasam K, & Morocz IA (2010) The benefit of symbols: monkeys show linear, human-like, accuracy when using symbols to represent scalar value. Animal cognition 13(5):711–719.

5. Boysen ST & Berntson GG (1995) Responses to quantity: perceptual versus cognitive mechanisms in chimpanzees (Pan troglodytes). Journal of Experimental Psychology: Animal Behavior Processes 21(1):82.

6. Fodor JA (1980) Fixation of belief and concept acquisition.

7. Frank MC, Slemmer JA, Marcus GF, & Johnson SP (2009) Information from multiple modalities helps 5-month-olds learn abstract rules. Developmental Science 12(4):504–509.

8. Johnson SP, et al. (2009) Abstract rule learning for visual sequences in 8-and 11-month-olds. Infancy 14(1):2–18.

9. Marcus GF, Fernandes KJ, & Johnson SP (2007) Infant rule learning facilitated by speech. Psychological Science 18(5):387–391.

10. Marcus GF, Vijayan S, Bandi Rao S, & Vishton PM (1999) Rule Learning by Seven-Month-Old Infants. Science 283(5398):77–80.

11. Gervain J, Macagno F, Cogoi S, Peña M, & Mehler J (2008) The neonate brain detects speech structure. Proceedings of the National Academy of Sciences 105(37):14222–14227.

12. Kouider S, et al. (2013) A neural marker of perceptual consciousness in infants. Science 340(6130):376–380.

13. Kouider S, et al. (2015) Neural dynamics of prediction and surprise in infants. Nature communications 6.

14. Emberson LL, Richards JE, & Aslin RN (2015) Top-down modulation in the infant brain: Learning-induced expectations rapidly affect the sensory cortex at 6 months. Proceedings of the National Academy of Sciences 122(31):9585–9590.

15. Norcia AM, Appelbaum LG, Ales JM, Cottereau BR, & Rossion B (2015) The steady-state visual evoked potential in vision research: a review. Journal of vision 15(6):4–4.

16. Walter W, Cooper R, Aldridge V, McCallum W, & Winter A (1964) Contingent negative variation: an electric sign of sensori-motor association and expectancy in the human brain. Nature 203:380–384.

17. Mento G & Valenza E (2016) Spatiotemporal neurodynamics of automatic temporal expectancy in 9-month old infants. Scientific reports 6.

18. Karmiloff-Smith A (1995) Beyond modularity: A developmental perspective on cognitive science (MIT press).

19. Namy LL (2001) What’s in a name when it isn’t a word? 17-month-olds’ mapping of nonverbal symbols to object categories. Infancy 2(1):73–86.

20. Sidman M, et al. (1982) A search for symmetry in the conditional discriminations of rhesus monkeys, baboons, and children. Journal of the Experimental Analysis of Behavior 37(1):23–44.

21. Lionello-DeNolf KM (2009) The search for symmetry: 25 years in review. Learning & behavior 37(2):188–203.

22. Needham A & Baillargeon R (2000) Infants’ use of featural and experiential information in segregating and individuating objects: a reply to. Cognition 74(3):255–284.

23. Sloutsky VM (2003) The role of similarity in the development of categorization. Trends in cognitive sciences 7(6):246–251.

24. Xu F (2002) The role of language in acquiring object kind concepts in infancy. Cognition 85(3):223–250.

25. Medam T, Marzouki Y, Montant M, & Fagot J (2016) Categorization does not promote symmetry in Guinea baboons (Papio papio). Animal cognition 19(5):987–998.

26. Meyer T & Olson CR (2011) Statistical learning of visual transitions in monkey inferotemporal cortex. Proceedings of the National Academy of Sciences 108(48):19401–19406.

27. Ogawa A, Yamazaki Y, Ueno K, Cheng K, & Iriki A (2010) Neural correlates of species-typical illogical cognitive bias in human inference. Journal of Cognitive Neuroscience 22(9):2120–2130.

28. Kojima T (1984) Generalization between productive use and receptive discrimination of names in an artificial visual language by a chimpanzee. International Journal of Primatology 5(2):161–182.

29. Matsuzawa T (1985) Use of numbers by a chimpanzee. Nature 315(6014):57–59.

30. Pepperberg IM & Pepperberg IM (2009) The Alex studies: cognitive and communicative abilities of grey parrots (Harvard University Press).

31. Dutoit T (1997) An introduction to text-to-speech synthesis (Springer).

32. Maris E & Oostenveld R (2007) Nonparametric statistical testing of EEG- and MEG-data. Journal of Neuroscience Methods 164(1):177–190.

33. Izard Vr, Dehaene-Lambertz G, & Dehaene S (2008) Distinct cerebral pathways for object identity and number in human infants. PLoS Biol 6(2):e11.

